# Epidermal Wnt signalling regulates transcriptome heterogeneity and proliferative fate in neighbouring cells

**DOI:** 10.1101/152637

**Authors:** Arsham Ghahramani, Giacomo Donati, Nicholas M. Luscombe, Fiona M. Watt

## Abstract

Canonical Wnt/beta-catenin signalling regulates self-renewal and lineage selection within the mouse epidermis. Although the transcriptional response of keratinocytes that receive a Wnt signal is well characterised, little is known about the mechanism by which keratinocytes in proximity to the Wnt-receiving cell are co-opted to undergo a change in cell fate. To address this, we performed single-cell mRNA-Seq on mouse keratinocytes co-cultured with and without the presence of beta-catenin activated neighbouring cells. We identified seven distinct cell states in cultures that had not been exposed to the beta-catenin stimulus and show that the stimulus redistributes wild type subpopulation proportions. Using temporal single-cell analysis we reconstruct the cell fate changes induced by neighbour Wnt activation. Gene expression heterogeneity was reduced in neighbouring cells and this effect was most dramatic for protein synthesis associated genes. The changes in gene expression were accompanied by a shift from a quiescent to a more proliferative stem cell state. By integrating imaging and reconstructed sequential gene expression changes during the state transition we identified transcription factors, including Smad4 and Bcl3, that were responsible for effecting the transition in a contact-dependent manner. Our data indicate that non cell-autonomous Wnt/beta-catenin signalling decreases transcriptional heterogeneity and further our understanding of how epidermal Wnt signalling orchestrates regeneration and self-renewal.

## 1. Introduction

The mammalian epidermis comprises interfollicular epidermis (IFE), hair follicles, sebaceous glands and sweat glands. Under steady-state conditions, each of these compartments is maintained by distinct populations of stem cells. However, following wounding each stem cell subpopulation exhibits the capacity to contribute to all differentiated lineages^1^. Recent single-cell gene-expression profiling of adult mouse epidermis identified multiple epidermal subpopulations^2^. Furthermore, in cultures of human and mouse keratinocytes there are three or more subpopulations with varying proliferative potential^3,4^.

One pathway that plays a key role in regulating stem cell renewal and lineage selection in mammalian epidermis is Wnt/beta-catenin signalling, which is an important regulator of epidermal maintenance, wound repair and tumorigenesis^5,6^. Gene-expression profiling has identified a number of signalling pathways that are regulated by cell-intrinsic activation of beta-catenin. Wnt signalling is indispensable for adult epidermal homeostasis; loss of beta-catenin in the IFE causes a defect in stem-cell activation, resulting in reduced basal layer proliferation and IFE thinning^7,8^ and loss of hair follicles. Conversely, transient activation of epidermal beta-catenin in the adult epidermis leads to expansion of the stem-cell compartment and results in the formation of ectopic hair follicles at the expense of the sebaceous glands and an increase in IFE thickness^9,10^.

There is good evidence that intrinsic beta-catenin activation in epidermal keratinocytes leads to effects on neighbouring epidermal cells. For example, in the mouse hair follicle, activated mutant beta-catenin cells can co-opt wild type cells to form a new hair growth through secretion of Wnt ligands^11^. This form of non-cell autonomous (NCA) activation suggests that autonomous Wnt signalling has the capability of changing neighbour cell fate. Although the mechanisms of autonomous Wnt activation are well described, it is unclear how NCA effects differ to cell intrinsic effects and how beta-catenin can simultaneously regulate self-renewal while changing the fate of neighbouring cells.

In this study we set out to analyse NCA signalling in wild type mouse keratinocytes that were co-cultured with keratinocytes in which beta-catenin was activated. This has revealed previously unknown heterogeneity of wild type mouse keratinocytes and elucidated the effect of Wnt signalling on neighbouring cell state and heterogeneity.

## 2. Results

### 2.1 Single-cell mRNA-seq analysis of basal epidermal stem cells

To explore the effects of non-cell autonomous Wnt signalling on epidermal cell state we sequenced the transcriptomes of single wild type murine keratinocytes co-cultured with cells expressing an inducible form of stabilised beta-catenin (K14ΔNβ-cateninER) in a ratio of 9:1. We compared cells cultured in the absence of 4-hydroxy-tamoxifen (4OHT) with cells treated for 24h with Tamoxifen to induce betacatenin. Cells were then disaggregated, loaded onto the C1 96-well microfluidic device (Fluidigm) and captured for sequencing. Owing to the single-cell capture method used, highly keratinized and terminally differentiated cells over 20 microns in diameter were excluded. We identified K14ΔNβ-cateninER cells by aligning reads to the transgene sequence and subsequently removed these cells from analysis (5 untreated cells and 7 activated cells). After quality control we retained 71 wild type control cells and 67 wild type cells exposed to Wnt signalling neighbours. We recorded a median of 641,000 reads per cell equating to 4,000-8,000 genes expressed per cell (transcripts per million, TPM > 1). Read alignment distribution was in line with other single-cell RNA-seq datasets with minimal ribosomal and intergenic reads (Figure S1).

To explore cell-state heterogeneity in wild type keratinocytes that had not been exposed to a neighbour in which beta-catenin was activated (untreated sample) we used reverse graph-embedding, a machine-learning technique. This enabled us to reconstruct a parsimonious tree connecting all observed epidermal cell states (DDRTree, Monocle 2)^12^. We applied the DDRTree algorithm to wild type cells using expressed genes (TPM > 1) after removing cell-cycle associated genes. We identified seven distinct groups of cells (Figure 1A) representing varying states of proliferation and differentiation. States 1 and 7 showed markedly low expression of S100 epidermal differentiation complex genes in comparison with the remaining subpopulations, indicating they represent transcriptomic states prior to commitment to differentiation (Figure 1B; upper) ^13^. Additionally, State 1 showed the highest expression of Itgb1, a marker of cells with high colony-forming capacity *in vitro* (Figure 1B; lower)^4^. Taking these expression markers together we considered cells in States 3, 4 and 6 as early commitment to differentiation. Almost all cells highly expressed basal interfollicular epidermis markers such as Krt14 and Col17a1 as defined by the top 5 IFE basal markers from Joost (Figure 1C)^2^. For each cell state we determined genes differentially expressed versus the remainder of the population and identified between 6 (State 3) and 62 (State 5) markers (Figure 1D, Table S1). We observed that the top five markers for each state were sufficient to distinguish the self-renewing States 1 and 7 from other states, but not sufficient to separate the early differentiation-committed States 3, 6 and 4 (Figure 1E).

**Figure 1.**
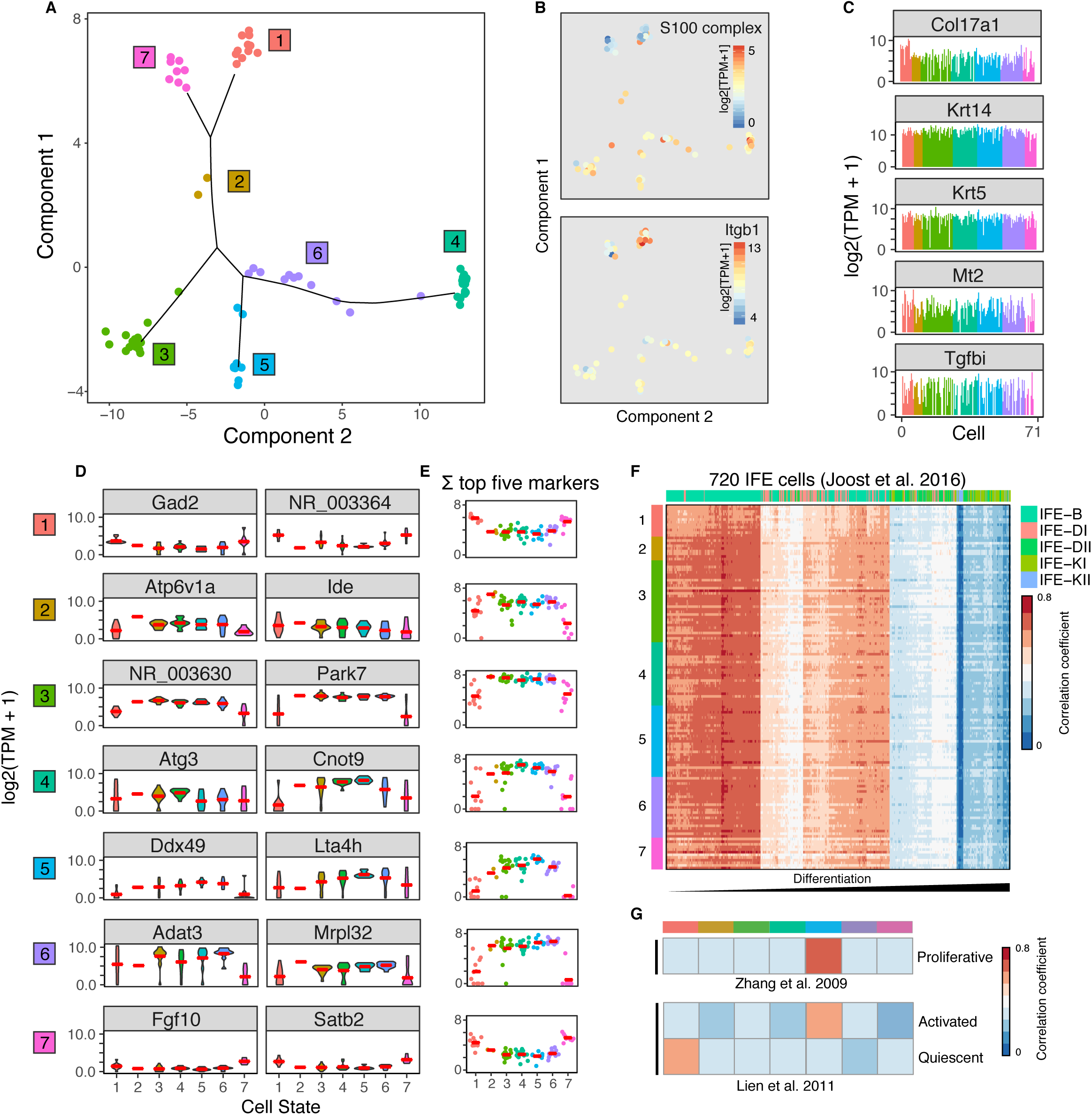
Molecular heterogeneity of epidermal cells in culture. **A)** Epidermal cell states classified by unsupervised clustering of single-cell transcriptomes. Data points represent 71 cells coloured according to 7 identified cellular states (labelled from 1 to 7). Axes represent the first two components of the dimensionally reduced transcriptomes. Lines depict the minimum spanning tree between the seven cell states. **B)** Upper: Expression of 17 S100 differentiation associated genes in each cell, displayed using the cell state map. Lower: Expression of Itgb1 in each cell. **C)** Expression levels of five basal interfollicular epidermal markers. Each bar represents one cell, bars are coloured by cell cluster. **D)** Distribution of the expression levels of cluster-specific markers coloured according to cell state. The top two differentially expressed markers are shown per cell state. Red line represents the median expression of the cluster. **E)** Summed expression levels of the top five differentially expressed markers for each state per cell. Red line presents the median per state. **F)** Heatmap showing the similarity between epidermal cells from this study and interfollicular epidermis (IFE) cells from Joost et al. Similarity is measured by Pearson’s correlation coefficient. Cells from this study are coloured by cluster along the vertical axis. Cells from Joost et al. are coloured by differentiation status along the horizontal axis. Joost et al. IFE cell legend is shown in order of differentiation status. B - basal IFE cells, DI and DII - differentiated suprabasal IFE cells, KI and KII - keratinized IFE cells. **G)** Heatmap showing similarity between average cluster transcriptomes from this study and proliferative IFE cells from Zhang et al. and activated versus quiescent IFE cells from Lien et al. 2011.

To determine the putative biological function of each cell state we correlated the single-cell signatures with a comprehensive set of IFE subpopulation expression profiles identified by Joost and colleagues. We further integrated two other bulk gene-expression studies which identified signatures for proliferative keratinocytes, activated and quiescent stem cells ^14,15^. From comparison with the Joost IFE subpopulations, all of our single cells correlate strongly with basal IFE stem cells, as expected since large (> 20μm) terminally differentiated cells were excluded from the analysis (Figure 1F). State 1 showed best correlation with an isolated subpopulation of quiescent self-renewing epidermal cells characterised by Lien and colleagues. (Figure 1G). State 1 and State 7 were highly correlated and shared many identified marker genes; however, they differed in their enrichment for an activated stem cell signature. State 7 showed moderate enrichment for activated stem-cell gene expression whereas State 1 showed no significant enrichment. We concluded that both are phenotypically self-renewing cells but differ in proliferative state. State 2 consisted of only two cells and therefore we did not characterise this subpopulation in detail; we hypothesise that it represents a transition state with lower stability than the other states, which would explain its low abundance. State 3 is a novel keratinocyte subpopulation: these cells could not be directly correlated with any previously studied *in vitro* or *in vivo* transcriptomic profiles. State 5 cells were characterized by expression of proliferation associated genes from Roshan (e.g. MRPL33, YY1) and correlated strongly with the expression profile of proliferative keratinocytes from Zhang^3,14^. Finally, States 4 and 6 were highly correlated with each other and DDRTree identified them as part of the same branch in the dimensionally reduced space. This branch of the state tree shows expression of early differentiation markers such as MXD1^16^ and highest expression of S100 early differentiation-associated genes. Figure 1A summarises the state classification of cells as determined by our cluster and DDRTree analysis highlighting the relationship between our seven identified keratinocyte states, depicted by branches.

### 2.2 Inducible Wnt signalling

In a previous study we generated gene-expression profiles from wild type and beta-catenin activated adult mouse epidermis^17^. We reanalysed these data to estimate the relative proportion of cells in each of the cell states identified *in vitro* (Figure 1A). We utilised CIBERSORT, a method for characterising the composition of tissue expression profiles resulting from mixtures of cells^18^. Our reanalysis indicated that epidermal beta-catenin signalling results in a depletion of cells in State 1 and increases the abundance of cells in State 5 (Figure 2A). This is consistent with the *in vivo* observation that intrinsic activation of epidermal beta-catenin results in proliferation and expansion of the stem cell compartment^10^ .

**Figure 2.**
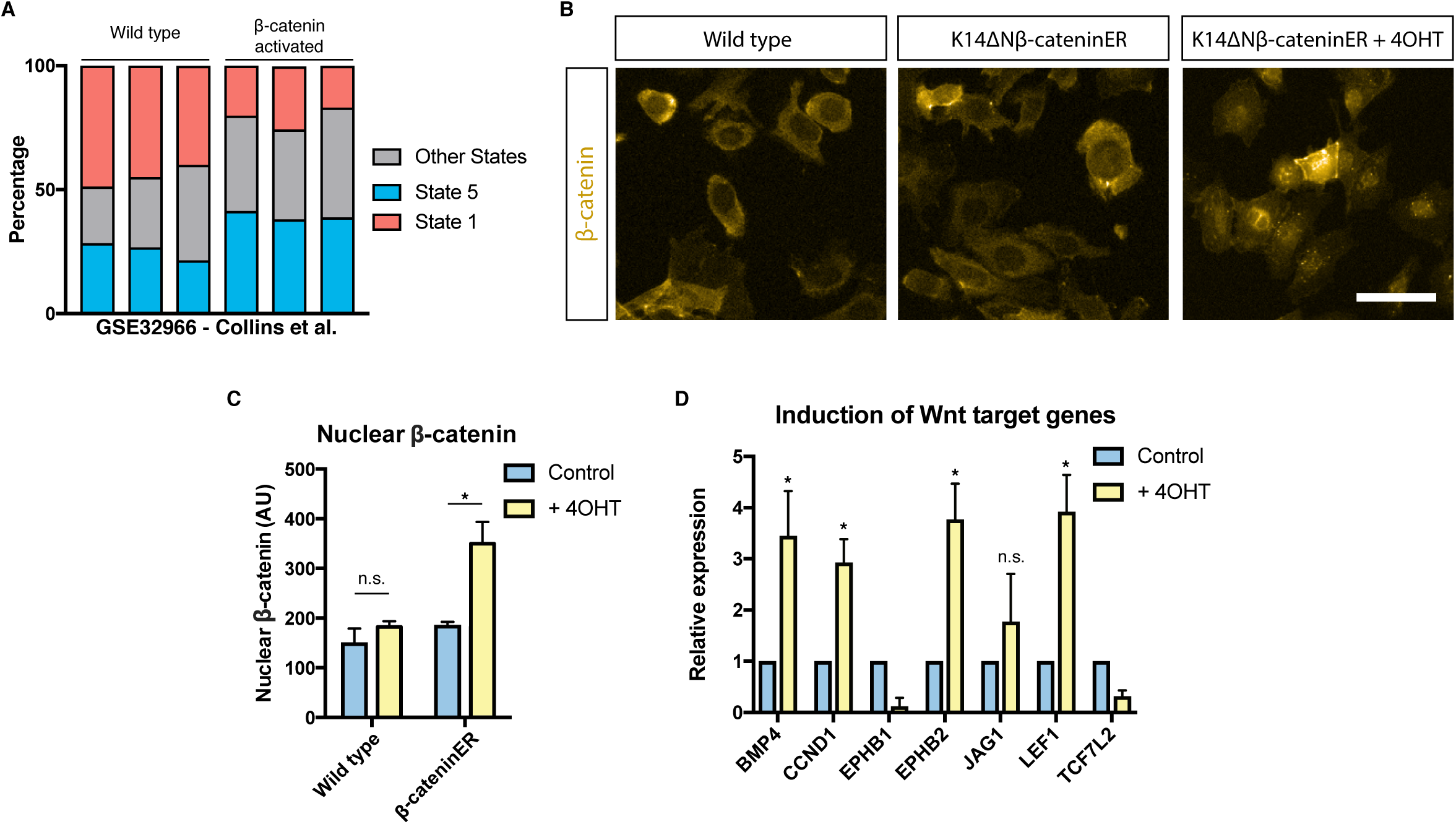
Induction of canonical Wnt signalling in a subpopulation of cells. **A)** Estimated proportions of State 1 and State 5 cells in wild type and beta-catenin activated epidermis samples from Collins et al. (GSE32966). **B)** Immunofluorescence showing cytoplasmic and nuclear beta-catenin in wild type and K14ΔNβ-cateninER keratinocytes after induction of canonical Wnt signalling using 4OHT. Scale bar, 20μm. **C)** Quantification of mean nuclear beta-catenin fluorescence intensity (n=3 independent cultures). **D)** Expression of canonical Wnt target genes in K14ΔNβ-cateninER cells upon induction with 4OHT quantified by RT-qPCR (n=4). *P < 0.05, n.s. Not significant. All data shown as mean ±SD.

Next to investigate whether epidermal cell states were altered by NCA Wnt signalling we examined the treated sample, wild type keratinocytes co-cultured with K14ΔNβ-cateninER keratinocytes in the presence 4OHT. We confirmed that K14ΔNβ-cateninER cells intrinsically activated canonical Wnt signalling in response to 4OHT by detecting beta-cateninER translocation into the nucleus (Figure 2B, C; Lo Celso et al., 2004). We also validated upregulation of canonical downstream target genes such as Bmp4, Cyclin-D1 and Lef1 in K14ΔNβ-cateninER keratinocytes using qRT-PCR (Figure 2D).

### 2.3 Reconstruction of NCA Wnt induced state transition

Having identified several different states of wild type keratinocytes and validated the intrinsic effects of beta-catenin activation, we used single-cell RNA-seq to deconvolve the effects of NCA Wnt signalling. Single-cell transcriptomes from wild type keratinocytes co-cultured with 4OHT-activated K14ΔNβcateninER cells were compared with those of wild type cells co-cultured with uninduced K14ΔNβcateninER cells and mapped onto the same dimensionally reduced space (Figure 3A). To exclude the possibility of transcriptional changes resulting from 4OHT treatment alone we screened bulk differential gene expression between the two cohorts of wild type cells. We found no evidence of estrogen receptor target genes among differentially expressed genes ^19^ .

**Figure 3.**
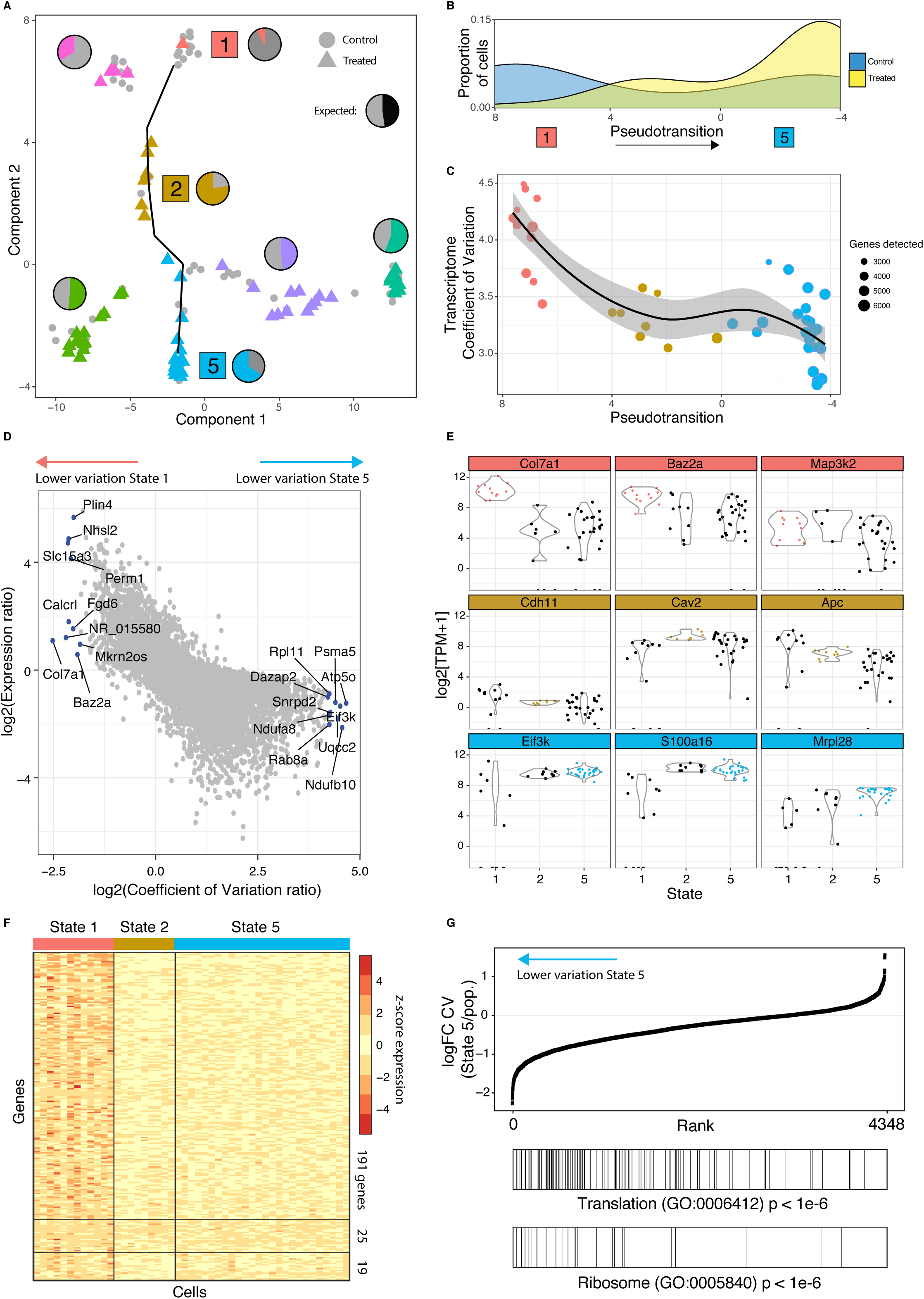
Neighbouring Wnt activation reduces gene expression heterogeneity. **A)** Keratinocytes exposed to Wnt signalling neighbours (NCA Wnt exposed) projected onto the wild type epidermal cell state map. States 1, 2 and 5 are labelled as these states contain different proportions of exposed versus unexposed cells. Wild type cells shown with grey circles (n = 71), exposed cells shown coloured by cell cluster as triangles (n = 71). Pie charts show relative proportions of wild type versus signalling-exposed cells in each cell state cluster. Line depicts the trajectory of cells changing from State 1 to State 5. **B)** Density of control versus NCA Wnt exposed cells along the reconstructed pseudotransition from States 1 to 5. **C)** Transcriptome coefficient of variation per cell (TCOV), coloured by cell state. Data point size represents number of genes detected as expressed (TPM > 1) for that cell. Line depicts a loess-curve fit for the pseudotransition-TCOV relationship. **D)** Scatterplot showing the log-ratio of coefficient of variation versus the log-ratio of gene expression between States 1 and 5. Labels for genes with the 10 highest and lowest coefficient of variation ratios. Also see Figure S2 for comparisons between State 1 and 2, and State 2 and 5. **E)** Violin plots depicting expression levels of cell state-specific genes showing differential dispersions between the three transition States 1, 2 and 5. **F)** Expression heatmap showing the reduction in gene expression heterogeneity of genes between States 1, 2 and 5 (z-score normalised log2(TPM + 1)). 191 genes show the most heterogeneous expression in State 1, 25 in State 2 and 19 in State 5. **G)** Gene rank enrichment analysis of change in gene coefficient of variation (CV) between cells in State 5 and all other cells. Genes with lower variation in state 5 are enriched for translation and ribosome-associated GO annotations (p-value < 1e-6).

We observed the same seven distinct transcriptional states in wild type cells with or without NCA Wnt signalling (Figure 3A), confirmed by independently analysing the treated cell population to reveal seven equivalent subpopulations. However, exposure to non-cell autonomous Wnt signalling markedly changed the state distribution of keratinocytes (Fisher’s exact test, p<0.05). Pie charts in Figure 3A show the observed ratio of control and signalling-exposed cells. States 1, 2 and 5 significantly deviated from the expected ratio (binomial test, p<0.05). After exposure to NCA signalling there was a depletion of cells in the self-renewing, quiescent State 1 and a higher than expected proportion of cells in States 2 and 5, representing a transition towards a proliferative and more differentiated transcriptional state (Figure 3A).

Taking the states with altered cell proportions (States 1, 2 and 5), we reconstructed the state transition induced by neighbouring Wnt+ keratinocytes using the Monocle pseudotime method ^12^. Wild type and exposed cells were ordered from State 1 to State 5 to reconstruct the temporal order of gene expression changes for cells undergoing this transition, referred to as the pseudotransition. Figure 3B shows the proportion of control and Wnt signalling exposed cells along the reconstructed temporal transition from State 1 to State 5; from this distribution it is clear that NCA Wnt exposed cells bias towards State 5.

We next sought to understand why State 1 cells were uniquely depleted after neighbour Wnt signalling. Previous studies have shown that cell responses to extrinsic signalling are affected by intracellular and intercellular transcriptional noise^20–22^. We thus hypothesised that the response to NCA Wnt signalling involves changes in both the dynamic range of transcriptional variation (intracellular variation) and state-specific gene expression (intercellular variation).

### 2.4 NCA Wnt signalling reduces heterogeneity in protein synthesis-associated transcripts

We first examined whether there was a difference in intracellular transcriptomic heterogeneity between the three altered states and whether changes occurred along the pseudotransition. The resulting ordering of cells from State 1 to State 5 was used to examine the transcriptome coefficient of variation (TCOV) per cell (Figure 3C). Here TCOV is an intracellular measure of the spread of transcript abundance accounting for mean abundance. Notably, TCOV decreased over the state transition and was statistically significantly lower in States 2 and 5. This reduction in dynamic range of gene expression is consistent with previous studies that have shown that progenitor cells have a higher rate of stochastic multilineage gene expression which reduces upon cell-fate commitment ^23,24^.

Next we contrasted the heterogeneity of genes that do not change in expression level between the three transition states (Figure S2A-C). Figure 3D displays the relationship for the log-ratio of intercellular gene expression variation and expression level between the two extremes of the pseudotransition, States 1 and 5, with the top 10 differentially dispersed genes labelled. Of interest are genes which change in expression heterogeneity from State 1 to State 5 while remaining at constant expression levels. Notably, Baz2a, Sox2, Col7a1 and Calcrl were amongst the genes with reduced COV in State 1 without significant differential expression (Supplementary Table S2). Baz2a has been previously established as part of the nucleolar remodelling complex that is important for establishing epigenetic silencing and transcriptional repression of rRNA genes^25,26^. Sox2 is an adult stem cell factor shown to be expressed in multiple epithelia ^27^. Sox2 has been previously reported to be expressed in hair follicles but absent from the interfollicular epidermis ^28^. Decreased Sox2 transcriptional heterogeneity in State 1 may reflect its multilineage progenitor identity *in vitro*. Similarly Col7a1 and Calcrl are significantly upregulated in hair follicle bulge stem cells^29^.

We observed many more heterogeneously expressed genes in State 1 (191 genes) than either States 2 or 5 and the contrast in number of differentially dispersed genes is demonstrated using the symmetric expression scale in Figure 3F. In comparison, we observed only 14 significantly differentially heterogeneous genes with lower heterogeneity in State 2. A striking number of these are known regulators of stem cell identity such as Cdh11, Cav2 and Apc ^30–32^. They are typically upregulated in basal stem cells; however little is known about how their heterogeneity affects cell fate. Our analysis of intercellular variation suggests that in slow–cycling stem cells (State 1) with low RNA and protein metabolism^33^ , transcriptional heterogeneity is lowest for stem-cell marker genes, emphasising the importance of transcriptional noise in addition to transcriptional amplitude. These observations are consistent with our hypothesis that State 1 cells are responsive to NCA Wnt signalling due to greater transcriptional variability. Exposure to this coordinated extracellular stimulus reduces transcriptional heterogeneity for these cells and biases their fate towards State 5.

To determine genes essential for a cell to be receptive to neighbour Wnt activation we analysed the fold-change in heterogeneity between States 1 and 5, comprising the majority of genes with differential heterogeneity. We found strong enrichment for translation and ribosome related genes, indicating a role for protein synthesis (p < 1e-6, Figure 3G). We hypothesise that cells in State 1 exhibit a multilineage primed transcriptional programme with stochastic expression of metabolism associated genes. Upon fate commitment, cells in the IFE steadily increase their translational rate in a proliferation independent manner ^33^. Hence translation associated genes are subject to greater transcriptional regulation post-commitment independent of transcription level.

These data and our single-cell analysis identified an NCA Wnt-receptive subpopulation, State 1, with greater dynamic range in gene expression (TCOV) and greater variation in the abundance of protein synthesis associated transcripts. Introduction of the NCA Wnt stimulus reduces variability in both aspects.

### 2.5 Transcription factors driving cell fate change

To understand drivers of the observed differential heterogeneity we reconstructed transcriptional changes over time along the state trajectory. Expression of each gene was modeled as a nonlinear function of pseudotransition time^12^. We found 436 genes that were dynamically regulated over the state transition (False discovery rate < 5%; Figure 4A). Using hierarchical clustering we grouped these genes into four patterns of dynamic expression. Group I genes, mostly highly expressed in State 1, were enriched for methylation associated genes and histone modifiers such as Setd3 and Kdm7a. These genes represent a pre-transition transcriptional profile of State 1 cells without exposure to signalling beta-catenin induced cells. Group II genes show highest expression in State 2, the intermediate transition state, with enrichment for desmosome genes such as Dsc2, Dsc3, Dsg2 and Dsp, which are most highly expressed in the suprabasal layers of murine epidermis indicative of early commitment to differentiation ^2^. Group III and IV genes, predominantly expressed in State 5, were enriched for protein synthesis associated genes and entry into mitosis, respectively. Notably group II includes the transcription factor Klf5, a regulator of proliferation in intestinal epithelial cells ^34^, and E2f1,which leads to epidermal hyperplasia when overexpressed in mice^35^ .

**Figure 4.**
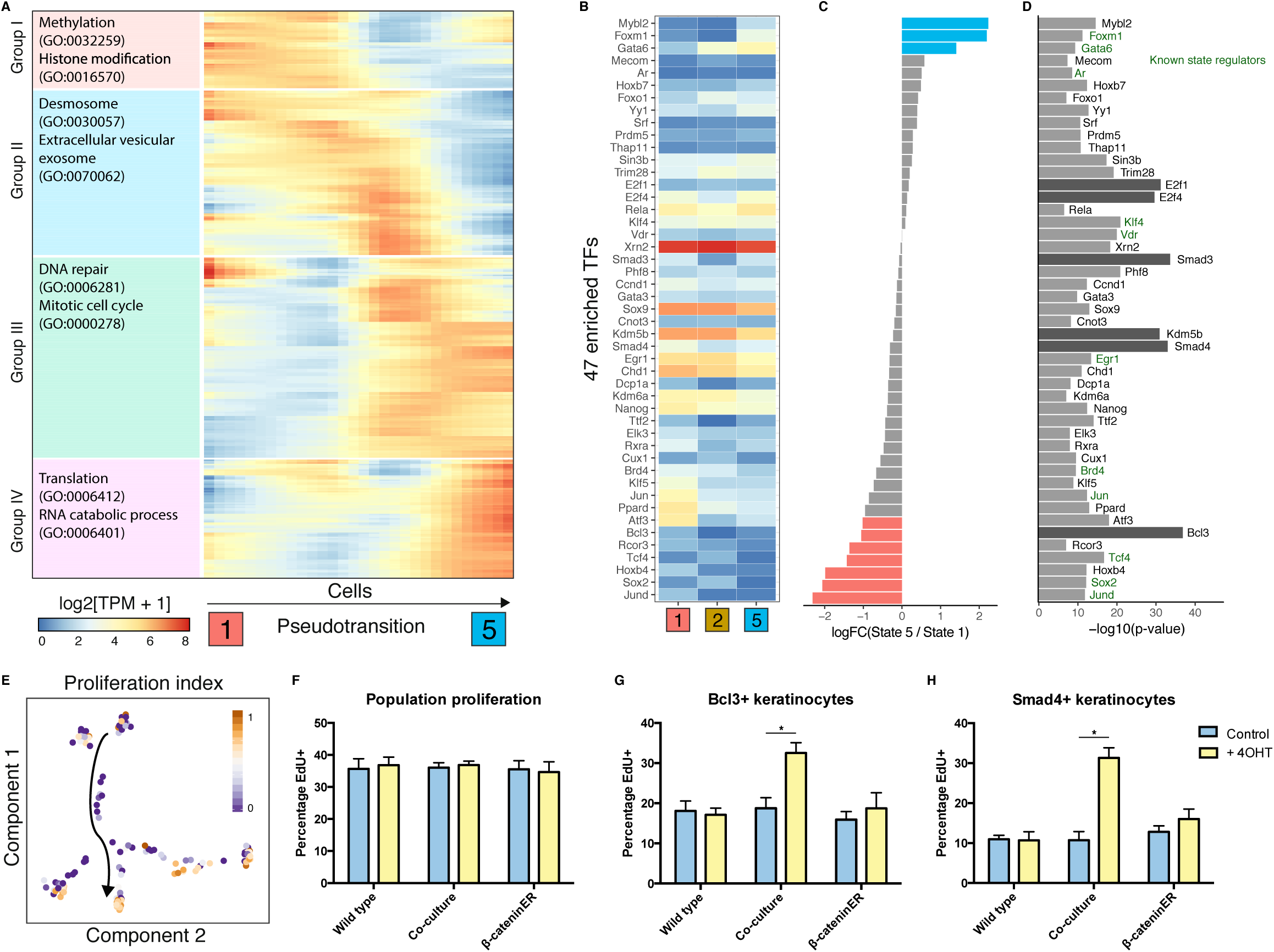
Induction of proliferation in Wnt+ neighbours is regulated by 47 TFs. **A)** Heatmap showing smoothed expression of pseudotransition-dependent genes (n = 436) ordered by hierarchical clustering and maximum expression. Top two enriched GO terms shown on left (all significant at q < 0.05). Genes (rows) are ordered by peak expression from State 1 to State 5. **B-D)** Heatmap showing expression of 47 transcription factors enriched as regulators of pseudotransition-dependent genes **(B)**. Log-fold change in expression of TFs between States 5 and 1 to show TFs with strong differential expression between the two states **(C)** and enrichment level of TFs **(D)**. **E)** Normalised proliferation index obtained by averaging expression of cell proliferation markers, projected onto the cell state map. Arrow denotes the direction of pseudotransition. **F-H)** Quantification of population proliferation by EdU assay in wild type, K14ΔNβ-cateninER and co-cultured keratinocytes with and without 4OHT treatment **(F)**. After stratification by Bcl3 nuclear abundance, cells in co-culture with activated K14ΔNβ-cateninER cells show a relative higher proliferation rate **(G)**. Similarly, stratification by nuclear Smad4 shows higher proliferation in the treated co-culture condition **(H)** (n = 3 independent cultures). *P < 0.05. All data shown as mean ±SD.

To gain insight into the regulation of the dynamically expressed genes induced by Wnt+ cells we performed a transcription factor motif analysis (Figure 3B-D). We calculated enrichment of transcription factor binding sites from the ChEA database, removing transcription factors (TFs) which were not expressed in any of the seven cell states (log[TPM+1] > 1). By analysing the promoters of the 436 dynamically expressed genes we identified 47 transcription factors putatively regulating the state transition. TFs were separated into three groups according to the directionality of gene expression from State 1 to 5: positive, negative and neutral. We noted that the activities of some identified TFs such as Smad3 and Smad4 are only partially dependent on expression level. Hence we did not rule out TFs on the basis of expression.

From this analysis we predicted Smad3, Smad4, Kdm5b, E2f1 and E2f4 as previously unknown key regulators of the state transition, with Bcl3 as a likely regulator of the specific transition between State 1 and 5. Known regulators of keratinocyte cell state are shown in blue (Figure 4D). Of note are Gata6 and Foxm1, two TFs upregulated in State 5 and previously shown to mark cells with multi-lineage differentiation potential and increased proliferative capacity respectively^36–38^ . We hypothesised that epidermal cells could be stratified based on the nuclear abundance of our identified novel state-regulating TFs, specifically Smad4 and Bcl3. Furthermore, our analysis on State 5 gene expression markers and transcriptome correlation indicated that this state shows a higher proliferation rate relative to States 1 and 2. To investigate further, we calculated a cell proliferation index consisting of normalised expression of three proliferation markers Pcna, Mcm2 and Mki67 (Figure 4E). This index demonstrated that State 5 comprises proliferative cells with few cells in States 1 or 2 expressing Pcna, Mcm2 or Mki67.

**Figure 5.**
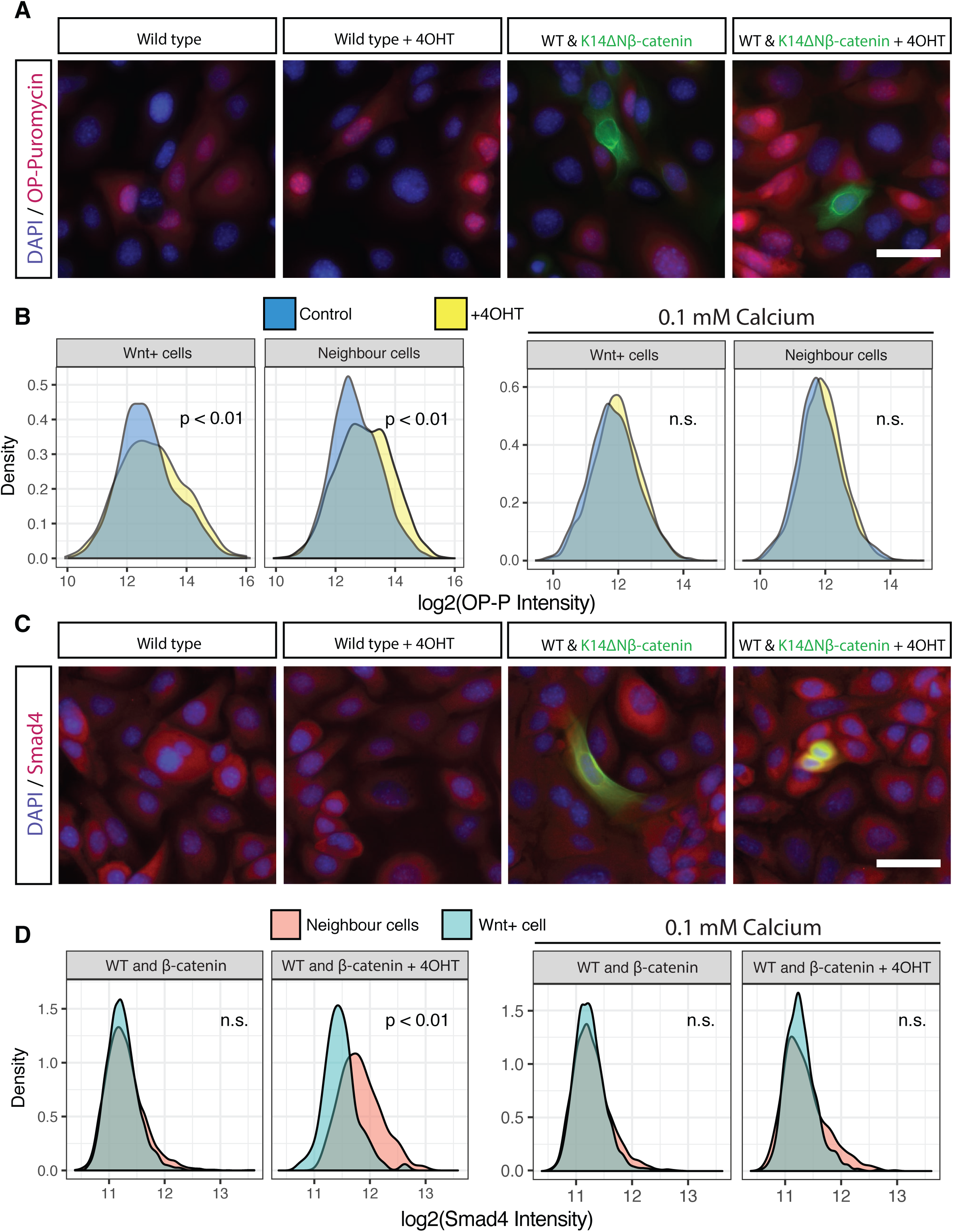
Wnt+ induced cell state transition is contact dependent. **A)** Appearance of typical wild type cells and K14ΔNβ-cateninER co-cultures stained for O-propargylpuromycin. **B)** Left: Quantification of O-propargyl-puromycin in K14ΔNβ-cateninER cells and their neighbours in control and 4OHT treated conditions. Right: Similar quantification for cells co-cultured in low calcium medium (0.1mM) to inhibit cell-cell contact (n=3 independent cultures, n=3 technical replicates, all pooled). **C)** Appearance of typical wild type cells and K14ΔNβ-cateninER co-cultures stained for Smad4. **D)** Left: Quantification of Smad4 in K14ΔNβ-cateninER cells and their neighbours in control and 4OHT treated conditions. Right: Similar quantification for cells co-cultured in low calcium medium (0.1mM) to inhibit cell-cell contact (n=3 independent cultures, n=3 technical replicates, all pooled). All scale bars, 20μm, n.s. Not significant.

To confirm our findings, we used an EdU incorporation assay to distinguish proliferating cells and analysed whether keratinocytes positive for our predicted driver TFs (measured by nuclear intensity) were more likely to be proliferative. At the population level there was no significant difference in proliferation when epidermal cells were co-cultured with 4OHT induced K14ΔNβ-cateninER cells (Figure 4F). However, when cells were discriminated by nuclear intensity for Bcl3 or Smad4 we observed a significant difference between wild type cells exposed to NCA Wnt signalling and wild type or induced K14ΔNβ-cateninER cell alone (Figure 4G and 4H). On average 18% of Bcl3^+^ cells were positive for EdU uptake in wild type or uninduced K14ΔNβ-cateninER cells; however, when wild type cells were co-cultured with induced K14ΔNβ-cateninER cells the EdU^+^ fraction rose to 34%. The proportion of proliferative Smad4^+^ cells increased from 10-18% to 33% EdU^+^.

Taken together these novel results indicate that Bcl3 and Smad4 are specific markers of the epidermal state transition and mark cells moving along the trajectory between States 1, 2 and 5 (Figure 1) during the first 24 hours of exposure to a NCA Wnt signal.

### 2.6 NCA Wnt induced state transition is contact dependent

From our total population of 67 cells exposed to NCA Wnt signalling, cells in States 3, 4 and 6 appeared to be unaffected. Our data suggest that State 1 comprises cells in a “responder” state permissive to NCA Wnt signalling due to the presence of key TFs and a more stochastic gene expression programme. We sought to address whether the reduction in ribosome-related gene expression heterogeneity and the induced expression of transition TFs are contact or distance dependent. To answer this question we labelled co-cultures of wild type and K14ΔNβ-cateninER cells with a cell reporter of protein synthesis.

We measured global protein translation by assaying incorporation of O-propargyl-puromycin assay (OPP) and compared wild type cells in contact with induced K14ΔNβ-cateninER cells to untreated control cells (Figure 5A). We observed that wild type cells showed higher translational activity when in contact with a Wnt+ cell. To confirm this we analysed the neighbours of over 10,000 Wnt+ cells and compared the OPP fluorescence intensity distributions (Figure 5B). We found a small but statistically significant increase in translation rate for both Wnt+ cells and neighbour cells in the 4OHT treated condition. This suggested a contact-dependent mechanism for control of protein synthesis downstream of NCA Wnt signalling.

To confirm contact dependence and to rule out local diffusion of soluble factors we repeated the assay in low calcium conditions. Keratinocytes cultured in low calcium medium do not form adherens or desmosomal cell contacts ^39,40^. Strikingly we observed no NCA Wnt effect under these conditions (Figure 5B). Similarly we observed no increase in nuclear abundance of Smad4, our predicted TF downstream of NCA Wnt signalling, in Wnt+ cells, as predicted. However, in neighbouring cells there was a significant increase in nuclear Smad4 intensity, which is abrogated in low calcium conditions (Figure 5C and 5D).

Taken together these data suggest that the Smad4-mediated cell state transition is downstream of non-cell autonomous Wnt signalling. Furthermore, the induction of this transition is contact dependent and does not occur under conditions where desmosomal adherens junctions are inhibited.

## 3. Discussion

Heterogeneity in the self-renewal and proliferative capabilities of keratinocytes has long been recognised. Previous analysis of clones and subclones of cultured human epidermal cells has demonstrated that there are at least three subpopulations, ‘holoclones’, ‘meroclones’ and ‘paraclones’ with descending self renewal potential^4,41,42^. More recently, Roshan et al. have shown the existence of two *in vitro* states with differing proliferation rates, and single cell transcriptomics have identified two distinct subpopulations of human keratinocytes in culture. In this study we have dissected molecular heterogeneity of epidermal cells at greater resolution and extended previous research by exploring the response of keratinocytes to neighbouring cells in which beta-catenin is activated. We have identified seven distinct transcriptomic states and characterised their biological relevance in order to create a state map of keratinocytes *in vitro*.

Using the cell state map and inducible activation, we have shown that Wnt/beta-catenin signalling acts to perturb cell fate by co-opting neighbours to become biased towards a pre-existing proliferative fate (Figure 6). It is important to note that we found no evidence for a *de novo* cell state as a result of non-cell autonomous signalling. This highlights the relevance of transient Wnt/beta-catenin signalling to cell state and is consistent with a model of stochastic epidermal commitment where extrinsic cues alter the likelihood of a cell switching state^43^. The observed difference in transcriptome variability between States 1 and 5 reflects a difference in cell state stability. Only a modest increase in translational activity is observed in State 5 or neighbouring cells; however, there is a marked reduction in the variability of translation-associated genes, highlighting the importance of transcriptional noise as well as transcriptional volume for determination of cell state.

**Figure 6.**
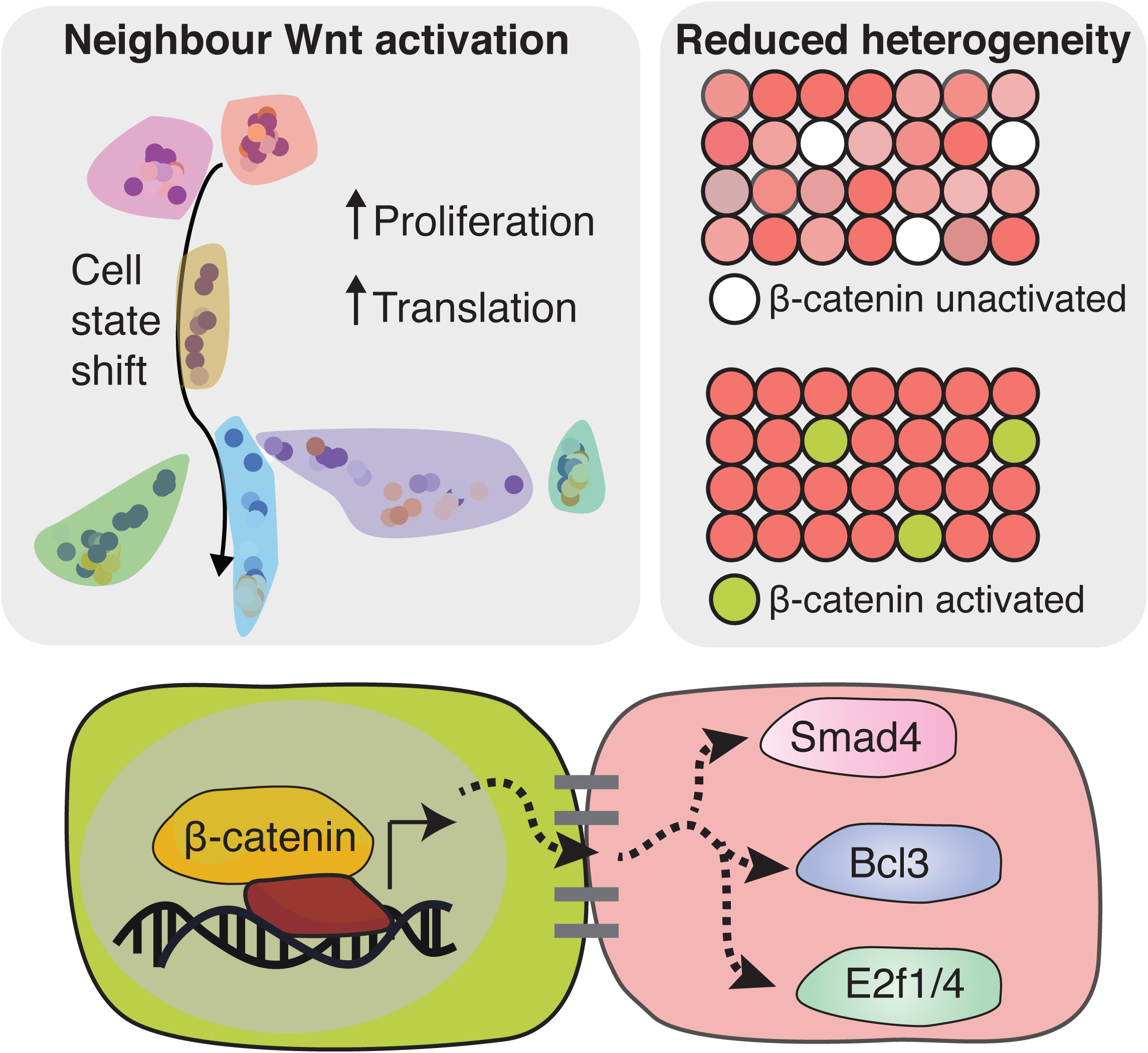
Graphical abstract - Wnt/beta-catenin activation affects neighbouring cells. Epidermal cells in culture adopt one of seven distinct transcriptomic states differing on the basis of proliferation and commitment to differentiation. Wnt/beta-catenin signalling acts as a non-cell autonomous signalling cue to activate a handful of transcription factors including Smad3/4, E2f1/4 and Bcl3 in neighbours. Concurrently, transcriptional heterogeneity is reduced as neighbour cells enter into a committed and proliferative transcriptional state.

Combined transcriptomic and positional single cell analyses allowed us to resolve spatial and temporal effects. As a result of this, we identified a collection of transcription factors, many of which were not previously implicated in epidermal cell state. One example is Bcl3, which is expressed in murine and human basal IFE, however, its role in epidermal cell fate is poorly understood^2,44^. In addition we identified Smad4 and utilised this as a marker of cell state transition. Smad4-beta-catenin cross-talk has been previously identified as essential for hair follicle maintenance ^45–47^. Here we show for the first time that beta-catenin signalling activation leads to Smad4 activation in a non-cell autonomous manner.

Our study does not address the extracellular effectors of NCA Wnt signalling. We previously identified a diverse array of secreted signalling molecules downstream of canonical Wnt signalling, including Bmp6, Dkk3, Wnt ligands, cytokines and ECM components^48^. Our analysis of TF effectors and previous evidence of epidermal self-renewal via autocrine Wnt signalling suggests that cells neighbouring a betacatenin^+^ cell are simultaneously committed to a state of lesser self-renewal and greater proliferative abilities. This is achieved via a combination of Bmp signalling (effected through Smad3/4; Figure 6) and neighbouring Wnt inhibition (Dkk3, Lim et al.). Intriguingly these effects are contact-dependent, hinting at yet-unknown mechanisms of locally restricting these signalling molecules or major signalling contributions from other membrane-bound factors. The observed difference in translation rate and proliferation in neighbouring cells demonstrates asymmetric coupling of cell fates, an essential component of epidermal homeostasis to ensure a balance of cell fates and epidermal metabolism.

In conclusion, our data provide a framework for studying cell state in the interfollicular epidermis and extend our understanding of functional heterogeneity and NCA signalling. Using this knowledge we demonstrate how Wnt/beta-catenin signalling, an orchestrator of regeneration, homeostasis and tumorigenesis in multiple tissues, influences neighbouring cell fate.

## 4. Methods

### Cell isolation and culture

K14ΔNβ-cateninER transgenic mice were generated as previously described ^10^. Keratinocytes were isolated and cultured from adult dorsal skin in FAD medium (one part Ham’s F12, three parts Dulbecco’s modified Eagle’s medium, 1.8x10^-4^ M adenine), supplemented with 10% foetal calf serum (FCS) and a cocktail of 0.5 μg/ml hydrocortisone, 5 μg/ml insulin, 1 x 10^-10^ M cholera enterotoxin and 10 ng/ml epidermal growth factor (HICE cocktail) ^49^. For the co-culture scRNA-seq experiment, wild type and K14ΔNβ-cateninER keratinocytes were cultured on 12 well plates in a ratio of 9:1 for a total of 200,000 cells per well and allowed to attach for 24 hours. Subsequently, cells were treated with 4-OHT (200nM) or DMSO as a control. After 24 hours of treatment cells were trypsinised and resuspended as a single cell suspension.

### Single cell capture, library preparation and RNA-sequencing

Single keratinocytes were captured on a medium-sized (10-17μm) microfluidic chip (C1, Fluidigm). Cells were assessed for viability (LIVE/DEAD assay, Life Technologies) and C1 capture sites were imaged by phase contrast to determine empty and doublet capture sites. Cells were loaded onto the chip at a concentration of 300 cells μl^-1^. Doublet or non-viable cells were excluded from later analysis. Cell lysis, reverse transcription, and cDNA amplification were performed on the C1 Single-Cell Auto Prep IFC, as per the manufacturer’s instructions. For cDNA synthesis the SMART-Seq v4 Ultra Low Input RNA Kit (Clontech) was used. Single cell Illumina NGS libraries were constructed with Nextera XT DNA Sample Prep kit (Illumina). Sequencing was performed on Illumina HiSeq4000 (Illumina) using 100bp paired-end reads.

### Bulk RNA extraction and real-time qPCR

Total RNA was purified with the RNeasy mini kit (Qiagen) with on-column DNaseI digestion, according to the manufacturer’s instructions. RNA was reverse transcribed with SuperScript III (Invitrogen). PCR reactions were performed with TaqMan Fast Universal PCR Master Mix and Taqman probes purchased from Invitrogen.

## RNA-seq quantification and statistical analysis

### Processing of reads and quality control

Reads were preprocessed using FastQC ^50^ and Cutadapt ^51^. Sequences were aligned to the Mus Musculus genome (GRCm38) using Tophat ^52^ discarding multiply-mapped reads. Gene level counts were extracted using featureCounts ^53^. Transcript levels were quantified as transcripts per million (TPM). Genes with a TPM greater than 1 were considered as expressed. We filtered cells for analyses on the basis of number of aligned reads (> 200,000), percentage of ribosomal reads (< 2%) and number of genes expressed (> 2000). 138 cells were taken forward for analysis.

### Identification of K14ΔNβ-cateninER cells

K14ΔNβ-cateninER cells were identified by aligning RNA-seq reads to the transgene locus using bowtie2 ^54^. Subsequently, cell identity was confirmed using qRT-PCR with Fast SYBR Green Master Mix (ThermoFisher Scientific) using the primers ATGCTGCTGGCTGGCTATGGTCAG (forward) and ATAGATCATGGGCGGTTCAGC (reverse) spanning the beta-catenin estrogen-receptor junction.

### Dimensionality reduction, cell state map and pseudotransition

We performed dimensionality reduction and constructed the principal graph representing transitions between all possible cell states using DDRTree from Monocle2 ^12^. We initially performed this analysis on wild type cells to determine the unperturbed cell state map. Subsequently we applied the DDRTree algorithm on the Wnt+ cell exposed group to confirm that we independently achieve a similar cell state map. We used all 138 cells for the final transcriptomic state map and differential expression to obtain cell state marker genes. Cell clusters obtained from Monocle were confirmed by a combination of dimensionality reduction of the cells using t-distributed stochastic neighbour embedding (tSNE) ^55^ and cluster identification with DBSCAN ^56^. Expression profiles from this study were correlated with expression profiles from Joost et al. (single cell RNA-seq, GSE67602), Zhang et al. (bulk microarray, GSE16516) and Lien et al. (bulk microarray, GSE31028) using pearson correlation coefficient.

### Heterogeneity analysis

Differential gene dispersion was performed using the Kolmogorov-Smirnov test after subtracting group mean expression from each group. Differentially dispersed genes were defined as q-value < 0.05. We filtered for genes with a coefficient of variation (CV) fold change > 2 between the state in question and the remainder of the population. Enrichment of gene log-fold change in heterogeneity was performed using a mean-rank gene set enrichment test on GO Biological Process terms as described previously ^57^.

### Pseudotransition gene expression and transcription factor enrichment

Pseudotransition cell ordering was determined by applying the Monocle pseudotime algorithm to the states with significantly different proportions of control and Wnt+ exposed cells (states 1, 2 and 5). Gene ontology enrichment was performed on the resulting clusters of temporal gene expression using enrichR ^58^. Transcription factor enrichment was performed by quantifying overrepresentation of target genes in the set of temporally regulated genes using the ChEA ChIP-X transcription factor binding database ^59^ .

## Immunofluorescence, imaging and neighbour cell quantification

### Immunofluorescence staining

The following antibodies were used: β-catenin (1:250, Sigma), Smad4 (1:250, Sigma), Bcl3 (1:250, Sigma). For EdU experiments (Molecular Probes; C10337), half of the cell culture medium was replaced with medium containing EdU for a final concentration of 10 μM EdU 30 minutes before fixation. Similarly, for OPP experiments (Molecular Probes; C10456) half of the cell culture medium was replaced 30 minutes before fixation with medium containing OPP for a final concentration of 20 μM OPP. Cultured cells were fixed with 4% PFA for 10 minutes followed by permeabilisation with 0.1% Triton X-100 for 10 minutes at room temperature. Cells were blocked for 1 hour at room temperature with 1% BSA in PBS. Primary antibody incubation was carried out for 90 minutes at room temperature. Samples were labelled with Alexa Fluor (488, 555, 647)- conjugated secondary antibodies for 1 hour at room temperature. Cells were imaged within 24 hours using an Operetta or Operetta CLS High-content Imaging System (PerkinElmer). Single cell cytoplasmic and nuclear fluorescence intensities were quantified with Harmony software (PerkinElmer) and analysed in R.

### Neighbour cell quantification

For neighbouring cell quantification K14ΔNβ-cateninER cells were labeled with CellTracker Green CMFDA dye (Molecular Probes) according to the manufacturer’s instructions. Single cell fluorescence intensity data and positional information were analysed in R. For each K14ΔNβ-cateninER CellTracker+ cell the mean fluorescence intensity of neighbouring cells was calculated. Neighbouring cells were defined as the nearest cell within 20μm (nucleus-to-nucleus distance). The mean number of neighbours was 5.4, as expected from a hexagonal packing model below confluence with mean cell diameter of 8μm. K14ΔNβ-cateninER cells were excluded if more than two neighbouring cells were also CellTracker+. Fluorescence intensity distributions from biological and technical replicates were pooled and contrasted between conditions using the non-parametric Kolmogorov-Smirnov test.

### Data Availability

Raw single-cell sequencing data and a table of processed and normalised read counts (TPM; transcripts per million) have been deposited in the Gene Expression Omnibus (GEO) under the accession GSE99989. Raw and processed high-content imaging data are available upon request.

### Author Contributions

AG designed and executed experiments, performed bioinformatic analyses and wrote the paper. GD conceived the project and supervised the experiments. NML and FMW supervised the project and co-wrote the paper.

## Acknowledgements

We are most grateful to Abdul Sesay and Leena Bhaw for their advice and technical assistance in performing single cell transcriptomics. AG is a King’s/Crick PhD student. This work was supported by the Francis Crick Institute which receives its core funding from Cancer Research UK (FC001110), the UK Medical Research Council (FC001110), and the Wellcome Trust (FC001110). This work was funded by grants to FMW from the Wellcome Trust and MRC. We also gratefully acknowledge funding from the Department of Health via the National Institute for Health Research comprehensive Biomedical Research Centre award to Guy’s & St Thomas’ National Health Service Foundation Trust in partnership with King’s College London and King’s College Hospital NHS Foundation Trust. We thank Victor Negri and Angela Pisco for helpful advice and discussions throughout the project.

The authors have no conflicts of interest to declare.

## Supplementary Data

**Supplementary Figure 1.**
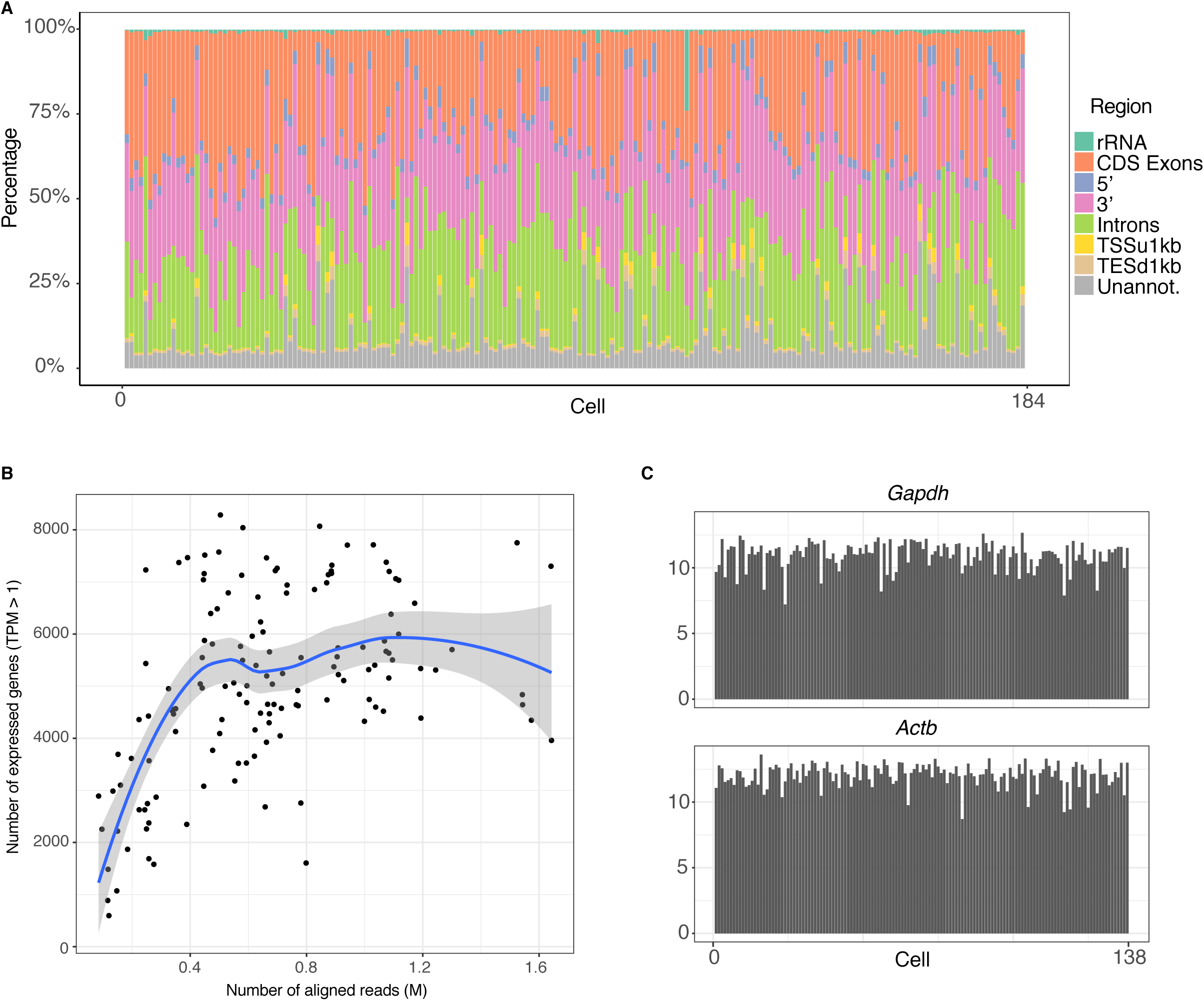
Quality control of single cell RNA-seq libraries. **A)** Stacked proportional barplot showing read alignment coverage distribution for all cells before QC filtering. TSSu1kb - region up to 1kb upstream of the transcription start site. TESd1kb - region up to 1kb downstream of the transcription end site. Unannot. - unannotated intergenic regions. **B)** Scatter plot showing relation between number of aligned reads and number of genes detected as expressed (TPM > 1). **C)** Barplots showing expression of two ubiquitously expressed genes (*Gapdh*, *Actb*) for 138 cells which have passed QC filtering.

**Supplementary Figure 2.**
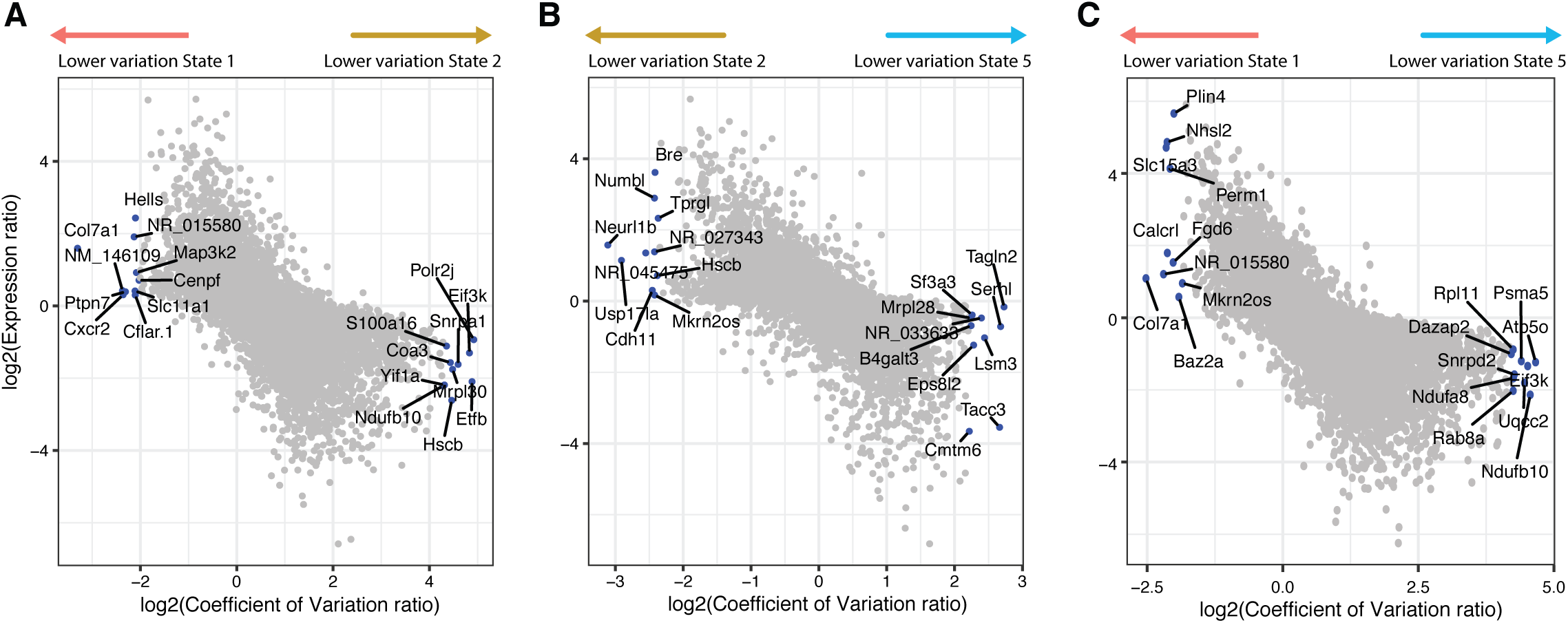
Comparison of heterogeneity and expression between transition states. A-C) Scatterplots showing the log-ratio of coefficient of variation versus the log-ratio of gene expression between pairs of pseudotransition states.

## Supplementary Tables

**Supplementary Table 1. Related to Figure 1E**

List of marker genes for each keratinocyte cell state (States 1-7).

**Supplementary Table 2. Related to Figure 3F**

Differentially dispersed genes along the NCA Wnt-induced cell state transition, i.e. in States 1, 2 and 5.

**Supplementary Table 3. Related to Figure 4A**

Genes dynamically expressed (and statistically significant) along the NCA Wnt-induced cell state transition. Cluster represents the four clusters of gene expression shown in Figure 4A.

**Supplementary Table 4. Related to Figure 4B-D**

List of transcription factors identified as regulating the NCA Wnt-induced cell state transition.

## References

1. Page, M. E., Lombard, P., Ng, F., Göttgens, B. & Jensen, K. B. The epidermis comprises autonomous compartments maintained by distinct stem cell populations. Cell Stem Cell 13, 471–482 (2013).

2. Joost, S. et al. Single-Cell Transcriptomics Reveals that Differentiation and Spatial Signatures Shape Epidermal and Hair Follicle Heterogeneity. cels 3, 221–237.e9 (2016).

3. Roshan, A. et al. Human keratinocytes have two interconvertible modes of proliferation. Nat. Cell Biol. 18, 145–156 (2016).

4. Jones, P. H. & Watt, F. M. Separation of human epidermal stem cells from transit amplifying cells on the basis of differences in integrin function and expression. Cell 73, 713–724 (1993).

5. Lim, X. & Nusse, R. Wnt signalling in skin development, homeostasis, and disease. Cold Spring Harb. Perspect. Biol. 5, (2013).

6. Watt, F. M. & Collins, C. A. Role of beta-catenin in epidermal stem cell expansion, lineage selection, and cancer. Cold Spring Harb. Symp. Quant. Biol. 73, 503–512 (2008).

7. Choi, Y. S. et al. Distinct functions for Wnt/β-catenin in hair follicle stem cell proliferation and survival and interfollicular epidermal homeostasis. Cell Stem Cell 13, 720–733 (2013).

8. Lim, X. et al. Interfollicular epidermal stem cells self-renew via autocrine Wnt signalling. Science 342, 1226–1230(2013).

9. Silva-Vargas, V. et al. β-Catenin and Hedgehog Signal Strength Can Specify Number and Location of Hair Follicles in Adult Epidermis without Recruitment of Bulge Stem Cells. Dev. Cell 9, 121–131 (2005/7).

10. Lo Celso, C., Prowse, D. M. & Watt, F. M. Transient activation of beta-catenin signalling in adult mouse epidermis is sufficient to induce new hair follicles but continuous activation is required to maintain hair follicle tumours. Development 131, 1787–1799 (2004).

11. Deschene, E. R. et al. β-Catenin activation regulates tissue growth non-cell autonomously in the hair stem cell niche. Science 343, 1353–1356 (2014).

12. Trapnell, C. et al. The dynamics and regulators of cell fate decisions are revealed by pseudotemporal ordering of single cells. Nat. Biotechnol. 32, 381–386 (2014).

13. Kypriotou, M., Huber, M. & Hohl, D. The human epidermal differentiation complex: cornified envelope precursors, S100 proteins and the ‘fused genes’ family. Exp. Dermatol. 21, 643–649 (2012).

14. Zhang, Y. V., Cheong, J., Ciapurin, N., McDermitt, D. J. & Tumbar, T. Distinct self-renewal and differentiation phases in the niche of infrequently dividing hair follicle stem cells. Cell Stem Cell 5, 267–278 (2009).

15. Lien, W.-H. et al. Genome-wide maps of histone modifications unwind in vivo chromatin states of the hair follicle lineage. Cell Stem Cell 9, 219–232 (2011).

16. Salehi-Tabar, R. et al. Vitamin D receptor as a master regulator of the c-MYC/MXD1 network. Proc. Natl. Acad. Sci. U. S. A. 109, 18827–18832 (2012).

17. Collins, C. A., Kretzschmar, K. & Watt, F. M. Reprogramming adult dermis to a neonatal state through epidermal activation of β-catenin. Development 138, 5189–5199 (2011).

18. Newman, A. M. et al. Robust enumeration of cell subsets from tissue expression profiles. Nat. Methods 12, 453–457 (2015).

19. Bourdeau, V. et al. Genome-wide identification of high-affinity estrogen response elements in human and mouse. Mol. Endocrinol. 18, 1411–1427 (2004).

20. Kolodziejczyk, A. A. et al. Single Cell RNA-Sequencing of Pluripotent States Unlocks Modular Transcriptional Variation. Cell Stem Cell 17, 471–485 (2015).

21. Guo, G. et al. Serum-Based Culture Conditions Provoke Gene Expression Variability in Mouse Embryonic Stem Cells as Revealed by Single-Cell Analysis. Cell Rep. 14, 956–965 (2016).

22. Shalek, A. K. et al. Single-cell RNA-seq reveals dynamic paracrine control of cellular variation. Nature 510, 363–369 (2014).

23. Hu, M. et al. Multilineage gene expression precedes commitment in the hemopoietic system. Genes Dev. 11, 774–785 (1997).

24. Velten, L. et al. Human haematopoietic stem cell lineage commitment is a continuous process. Nat. Cell Biol. 19, 271–281 (2017).

25. Gu, L. et al. BAZ2A (TIP5) is involved in epigenetic alterations in prostate cancer and its overexpression predicts disease recurrence. Nat. Genet. 47, 22–30 (2015).

26. Santoro, R., Li, J. & Grummt, I. The nucleolar remodeling complex NoRC mediates heterochromatin formation and silencing of ribosomal gene transcription. Nat. Genet. 32, 393–396 (2002).

27. Arnold, K. et al. Sox2(+) adult stem and progenitor cells are important for tissue regeneration and survival of mice. Cell Stem Cell 9, 317–329 (2011).

28. Driskell, R. R., Giangreco, A., Jensen, K. B., Mulder, K. W. & Watt, F. M. Sox2-positive dermal papilla cells specify hair follicle type in mammalian epidermis. Development 136, 2815–2823 (2009).

29. Blanpain, C., Lowry, W. E., Geoghegan, A., Polak, L. & Fuchs, E. Self-renewal, multipotency, and the existence of two cell populations within an epithelial stem cell niche. Cell 118, 635–648 (2004).

30. Kim, N.-H., Choi, S.-H., Lee, T. R., Lee, C.-H. & Lee, A.-Y. Cadherin 11, a miR-675 target, induces N-cadherin expression and epithelial-mesenchymal transition in melasma. J. Invest. Dermatol. 134, 2967–2976 (2014).

31. Tan, D. W. M. et al. Single-cell gene expression profiling reveals functional heterogeneity of undifferentiated human epidermal cells. Development 140, 1433–1444 (2013).

32. Chan, E., Gat, U., McNiff, J. M. & Fuchs, E. A common human skin tumour is caused by activating mutations in β-catenin. Nat. Genet. 21, 410–413 (1999).

33. Blanco, S. et al. Stem cell function and stress response are controlled by protein synthesis. Nature 534, 335–340 (2016).

34. Chanchevalap, S., Nandan, M. O., Merlin, D. & Yang, V. W. All-trans retinoic acid inhibits proliferation of intestinal epithelial cells by inhibiting expression of the gene encoding Krüppel-like factor 5. FEBS Lett. 578, 99–105 (2004).

35. Pierce, A. M. et al. Increased E2F1 activity induces skin tumors in mice heterozygous and nullizygous for p53. Proc. Natl. Acad. Sci. U. S. A. 95, 8858–8863 (1998).

36. Donati, G. et al. Wounding induces dedifferentiation of epidermal Gata6(+) cells and acquisition of stem cell properties. Nat. Cell Biol. 19, 603–613 (2017).

37. Gemenetzidis, E. et al. Induction of human epithelial stem/progenitor expansion by FOXM1. Cancer Res. 70, 9515–9526 (2010).

38. Molinuevo, R. et al. FOXM1 allows human keratinocytes to bypass the oncogene-induced differentiation checkpoint in response to gain of MYC or loss of p53. Oncogene 36, 956–965 (2017).

39. Hennings, H. & Holbrook, K. A. Calcium regulation of cell-cell contact and differentiation of epidermal cells in culture. An ultrastructural study. Exp. Cell Res. 143, 127–142 (1983).

40. O’Keefe, E. J., Briggaman, R. A. & Herman, B. Calcium-induced assembly of adherens junctions in keratinocytes. J. Cell Biol. 105, 807–817 (1987).

41. Barrandon, Y. & Green, H. Three clonal types of keratinocyte with different capacities for multiplication. Proc. Natl. Acad. Sci. U. S. A. 84, 2302–2306 (1987).

42. Jones, P. H., Harper, S. & Watt, F. M. Stem cell patterning and fate in human epidermis. Cell 80, 83–93 (1995).

43. Rompolas, P. et al. Spatiotemporal coordination of stem cell commitment during epidermal homeostasis. Science 352, 1471–1474 (2016).

44. Uhlén, M. et al. Proteomics. Tissue-based map of the human proteome. Science 347, 1260419 (2015).

45. Owens, P. et al. Smad4-dependent desmoglein-4 expression contributes to hair follicle integrity. Dev. Biol. 322, 156–166 (2008).

46. Qiao, W. et al. Hair follicle defects and squamous cell carcinoma formation in Smad4 conditional knockout mouse skin. Oncogene 25, 207–217 (2006).

47. Yang, L., Wang, L. & Yang, X. Disruption of Smad4 in mouse epidermis leads to depletion of follicle stem cells. Mol. Biol. Cell 20, 882–890 (2009).

48. Donati, G. et al. Epidermal Wnt/β-catenin signalling regulates adipocyte differentiation via secretion of adipogenic factors. Proc. Natl. Acad. Sci. U. S. A. 111, E1501–9 (2014).

49. Watt, F. M., Simon, B. & Prowse, D. M. in Cell Biology 133–138 (2006).

50. Babraham Bioinformatics - FastQC A Quality Control tool for High Throughput Sequence Data. Available at: http://www.bioinformatics.babraham.ac.uk/projects/fastqc/. (Accessed: 18th April 2017)

51. Martin, M. Cutadapt removes adapter sequences from high-throughput sequencing reads. EMBnet.journal 17, 10–12 (2011).

52. Kim, D. et al. TopHat2: accurate alignment of transcriptomes in the presence of insertions, deletions and gene fusions. Genome Biol. 14, R36 (2013).

53. Liao, Y., Smyth, G. K. & Shi, W. featureCounts: an efficient general purpose program for assigning sequence reads to genomic features. Bioinformatics 30, 923–930 (2014).

54. Langmead, B. & Salzberg, S. L. Fast gapped-read alignment with Bowtie 2. Nat. Methods 9, 357–359 (2012).

55. Maaten, L. van der & Hinton, G. Visualizing Data using t-SNE. J. Mach. Learn. Res. 9, 2579–2605 (2008).

56. Ester, M., Kriegel, H. P., Sander, J. & Xu, X. A density-based algorithm for discovering clusters in large spatial databases with noise. KDD (1996).

57. Michaud, J. et al. Integrative analysis of RUNX1 downstream pathways and target genes. BMC Genomics 9, 363 (2008).

58. Kuleshov, M. V. et al. Enrichr: a comprehensive gene set enrichment analysis web server 2016 update. Nucleic Acids Res. 44, W90–7 (2016).

59. Lachmann, A. et al. ChEA: transcription factor regulation inferred from integrating genome-wide ChIP-X experiments. Bioinformatics 26, 2438–2444 (2010).

